# Sleep Fragmentation Modulates the Neurophysiological Correlates of Cognitive Fatigue

**DOI:** 10.1101/2024.07.23.604738

**Authors:** Oumaïma Benkirane, Peter Simor, Olivier Mairesse, Philippe Peigneux

## Abstract

Cognitive fatigue (CF) is a critical factor affecting performance and well-being. It can be altered in suboptimal sleep quality conditions, e.g., in patients suffering from obstructive sleep apnea who experience both intermittent hypoxia and sleep fragmentation (SF). Understanding the neurophysiological basis of SF in healthy individuals can provide insights to improve cognitive functioning in disrupted sleep conditions. In this electroencephalographical (EEG) study, we investigated in 16 healthy young participants the impact of experimentally induced SF on the neurophysiological correlates of CF measured before, during, and after practice on the TloadDback, a working memory task tailored to each individual’s maximal cognitive resources. Participants spent two times three consecutive nights in the laboratory, once in an undisrupted sleep (UdS) condition and once in a SF condition induced by non-awakening auditory stimulations, counterbalanced, and performed the TloadDback task both in a high (HCL) and a low (LCL) cognitive load condition. EEG activity was recorded during wakefulness in the 5-minutes resting state immediately before and after, as well as during the 16-minutes of the TloadDback task practice. In the high cognitive load under sleep fragmentation (HCL/SF) condition, high beta power increased during the TloadDback indicating heightened cognitive effort, and beta and alpha power increased in the post- vs. pre task resting state, suggesting a relaxation rebound. In the low cognitive load/undisturbed sleep (LCL/UdS) condition, low beta activity increased suggesting a relaxed focus, as well as mid beta activity associated with active thinking. These findings highlight the dynamic impact of SF on the neurophysiological correlates of CF and underscore the importance of sleep quality and continuity to maintain optimal cognitive functioning.

## 1. Introduction

Continuous and high-quality sleep during the night is crucial for optimal brain and body functioning (Xie et al., 2013; Jiang et al., 2018; Laharnar et al., 2020). Conversely, sleep deprivation (SD) and/or poor sleep quality can have detrimental effects on both physical health and cognitive performance (Killgore, 2010; Sharma & Kavuru., 2010). While there is considerable variation in individual sleep requirements (Hirshkowitz et al., 2015), it is generally recommended that adults get an average 7 hours of sleep per night to avoid negative health consequences (Watson et al., 2015). However, even individuals with normal sleep duration may not experience efficient sleep (Touzet, 2017), potentially due to various factors impacting sleep quality and continuity (Ohayon, 2007). One factor that may affect sleep continuity is sleep fragmentation (SF), characterized by repeated interruptions during sleep that, even if not resulting in complete awakenings, prevent the brain from entering consolidated, slow-wave (Philip et al., 1994) and rapid eye movement (REM) sleep. SF of these stages was shown to impair the restorative effects of sleep (Durmer, & Dinges, 2005; Riemann, et al., 2012; Ren et al., 2023) and exert detrimental influence on daytime cognitive functions (Stepanski, 2002; Nair et al., 2011). Amongst others, SF is a prominent feature in sleep-related breathing disorders, particularly Obstructive Sleep Apnea (OSA), characterized by recurrent episodes of airflow obstruction leading to brief arousals, intermittent hypoxemia, snoring, and SF (Mannarino et al., 2012). Although studies demonstrated a specific impact of SF on cognitive performance in individuals with OSA (Alomri et al., 2021), there is still an ongoing debate regarding the respective effects of hypoxemia and SF on the ensuing daytime neurocognitive deficits (Alomri et al., 2021; Colt et al, 1991; Valencia-Flores et al., 2016). Nonetheless, SF is certainly a contributing factor to OSA-related disturbances, neurocognitive deficits being commonly reported in individuals with OSA in whom the restorative effects of sleep are impaired (Verstraeten, 2007; Krysta et al., 2017). Neurocognitive deficits predominantly impact vigilance (Daurat et al., 2016), but also performance in tasks that require continuous cognitive engagement, such as sustained attention and executive functions, and can also exert adverse effects on various aspects of memory, productivity, and social interactions (Csábi, 2013; Daurat et al., 2008; Djonlagic et al., 2012; Olaithe et al., 2013; Weaver & George, 2011). In healthy individuals, SF induced by noise was shown to increase upper airway collapsibility and heart rate, that are characteristic features of OSA (Sériès, 1994; Griefahn et al., 2008). Since the independent contributions of SF and hypoxemia to cognitive deficits are hard to disentangle in pathological conditions such as OSA (Alomri et al., 2021), studying the consequences of experimentally induced SF in healthy individuals may help better understanding the impact of disrupted restorative sleep mechanisms on cognitive processes and their underlying neurophysiological basis. In other terms, systematically examining the effects of experimentally induced and reversible SF in a healthy population may provide a valuable model to understand how and to what extent SF contributes to cognitive deficits associated with OSA.

One cognitive symptom both associated with SF and sleep-related breathing disorders is cognitive fatigue (CF), subjectively experienced as a decrease in cognitive efficiency in situations involving sustained cognitive demands and constrained processing time, independently of sleepiness (Mairesse et al., 2019). CF poses a significant concern as it can lead to errors, impaired performance in daily activities, decreased efficiency, motivation loss, and an increased risk of accidents (Fithriyyah, et al., 2020). The terminology and definitions used to describe CF widely vary throughout the literature (Hamann & Carstengerdes, 2023). One of the most notable definitions comes from Grandjean (1979), who describes mental fatigue as a state of reduced alertness that approaches, but does not reach, sleepiness. This state is induced by prolonged task execution and/or highly demanding tasks and can be mitigated by taking breaks. In this study, we adhere to Grandjean’s (1979) definition, which characterizes CF as acute and non-pathological, induced by demanding or prolonged tasks or when processing time is limited in mentally demanding activities, leading to subjective feelings of fatigue, tiredness, reduced energy, and decreased performance (Marcora et al., 2009). Symptoms of cognitive fatigue include difficulties concentrating (Holtzer et al., 2011), memory issues (Massar, et al., 2010), and decline in executive functions (Kato et al., 2009). Thus, CF distinguishes from sleepiness, or drowsiness, which is characterized by a strong urge to sleep or a tendency to fall asleep, usually due to insufficient sleep, disrupted sleep patterns, or certain medical conditions (Neu et al., 2010). Physical fatigue, on the other hand, involves a sensation of physical exhaustion and decreased physical performance caused by prolonged physical activity, lack of rest, or medical conditions affecting physical stamina and energy levels (Stone et al., 1987). While cognitive fatigue, sleepiness, and physical fatigue are interrelated, they can occur independently or together, depending on factors such as lifestyle, mental or physical health status, and external stressors. Different countermeasures are typically needed to alleviate each condition: active or passive rest for cognitive and physical fatigue, and sleep for sleepiness. However, sleep might also alleviate cognitive fatigue levels, although it may not always be effective, such as in cases of insomnia or chronic fatigue conditions (Neu et al., 2011).

Previous studies on CF mainly used tasks performed on extended periods even up to several hours with a constant level of cognitive demands, such as e.g., mental arithmetic calculations (Trejo et al., 2015; Lorist et al., 2000; Lorist et al., 2005; Mizuno et al., 2011; Van der Linden et al., 2003; Ackerman & Kanfer; 2009; Lim et al. 2010). Hence, these studies examined the impact of high cognitive task demands on CF, with the underlying assumption that sustained cognitive demands will eventually deplete cognitive resources and increase CF levels (Cook et al., 2007; Shigara et al., 2013). Other studies investigated neural compensation within a fatigue framework, wherein participants engage in a taxing cognitive task for an extended duration (Wang et al., 2014), consistently triggering cognitive fatigue. Among young adults, as cognitive fatigue intensifies, behavioral performance decreases, with observable signs of neural compensation mitigating fatigue-induced cognitive deficits (Wang et al., 2016). Recently, Borragán et al. (2017) proposed that CF can also be understood within the framework of the time-based resource-sharing (TBRS) model (Barrouillet and Camos, 2012). According to this model, it is not the complexity of the task itself, but the time allotted for processing incoming stimuli that determines cognitive load and, subsequently, fatigue. In this framework, attention is viewed as a finite resource that varies among individuals, and an individual’s maximal cognitive load corresponds to the fastest pace at which he or she can still accurately meet task demands. Working continuously at maximal cognitive load will eventually result in increased CF. Modulation of CF by cognitive load level (Borragán et al., 2017) as well as by sleep deprivation (Borragán et al., 2019), duration (Borragán et al., 2018) and fragmentation (Benkirane et al. 2022) have already been investigated using the TloadDback task (Borragán et al. 2017), a cognitive load task designed to account for interindividual variability in working memory processing capabilities.

However, the neurophysiological mechanisms underlying CF and its interactions with prior sleep quality and/or continuity, as well as their combined impact on performance in several cognitive domains remain poorly investigated. Most cognitive fatigue research in neuroscience has concentrated on examining changes in EEG patterns (Li et al., 2020), which are considered a promising biomarker of mental fatigue (Tran et al., 2020). Evidence supports the notion that mental fatigue induces discernible alterations in EEG signals, as estimated through power spectral density (PSD) (Tian et al., 2018). Within PSD, Individual Alpha Frequency emerges as a significant metric. Situated within the alpha frequency range (7.5 - 12.5 Hz), it is indicative of arousal, attention, and cognitive performance. It serves as a stable neurophysiological marker and aids in detecting physical fatigue (Fithriyyah, et al., 2020). CF has also been found to be associated with changes in frontal theta activity, a reliable correlate of cognitive control (Wascher, 2014). Alterations in resting-state EEG power induced by mental fatigue may offer valuable insights into identifying its neural mechanisms (Tanaka et al., 2012). Given the evidence that EEG changes serve as reliable indicators for detecting mental fatigue, EEG has emerged as one of the foremost technical methods to investigate the neurobiological correlates and detection of mental fatigue (Li et al., 2020).

In this context, we investigated using electroencephalography (EEG) in healthy young adults how experimentally induced SF exerts an impact on the neurophysiological correlates of CF at rest before and after, and during practice on the TloadDback working memory task, that allows tailoring cognitive load to each individual’s maximal capabilities. Participants spend two times three consecutive nights in the laboratory, once in a SF condition induced by non-awakening auditory stimulations, and once in a restorative, undisrupted sleep (UdS) condition, counterbalanced.

## 2. Materials and Methods

### 2.1 Participants

Sixteen young healthy participants (8 females; average 28.5 ± 4.48 years, range 24-38 years) were recruited through social media advertisements and flyers, and provided written consent to take part in this study approved by the Faculty and ULB-Erasme Hospital ethics committees (CE 001/2019). To avoid desirability bias, they were not informed of the exact purpose of the experiment and received a complete debriefing at the end of the study. Sleep or breathing disorders, irregular sleep patterns, extreme morning or evening chronotypes, habitual sleep duration lesser than 6.5 hours, neurological or psychiatric conditions, or a history of opioid treatment or current use of benzodiazepines were exclusion criteria. They were also instructed to refrain from consuming stimulating or alcoholic beverages on the day prior to the experimental days and during the experimental days.

### 2.2 General experimental procedure

The behavioral component of CF in this experiment is reported elsewhere (Benkirane, et al., 2022), to which the reader is referred for a detailed presentation. Only essential information is reported here. The study spanned a duration of 17 days, as depicted Figure 1. To confirm the absence of sleep or breathing disorders to facilitate habituation to the sleep lab conditions, participants were asked to spend the first night (Day 1) in the sleep laboratory under polysomnographic recordings (PSG) one week before the first experimental session. Regularity of the sleep-wake pattern was controlled using both daily sleep diary and visual inspection of wrist-worn actigraphy (wGT3XBT Monitor, ActiGraph, Pensacola, FL, USA). On day 8, participants went to the sleep laboratory for three consecutive nights under PSG recording, either in undisturbed sleep (UdS) or a sleep fragmentation (SF) condition. After three nights of sleep at home, participants returned for three consecutive PSG nights in the other condition (SF or UdS). The order of UdS and SF conditions was counterbalanced across participants. On SF and UdS experimental nights, participants arrived at the laboratory between 21:30 and 22:00 for PSG prepping. Lights were turned off between 23:00 and 24:00 depending on individual sleep habits, as well as their habitual sleep duration (mean 531.43 ± 58.82 minutes). In the SF condition, auditory stimuli were presented during the three consecutive nights to induce sleep fragmentation (see below SF procedure), while in the UdS condition, participants were allowed to sleep undisturbed for three consecutive nights. The morning following each UdS and SF night (days 9 to 11 and 15 to 17), they completed the St Mary’s Sleep Questionnaire (Ellis et al., 1981) to subjectively assess the quality of their previous night of sleep. Following the initial nights of undisturbed sleep (UdS) and sleep fragmentation (SF) on days 9 and 15, participants underwent a neuropsychological assessment battery and the calibration segment of the TloadDBack task, aimed at determining their peak working memory processing capacity (see below) (Borragán et al., 2017). Subsequently, during days 10–11 and 16–17, after the second and third UdS and SF nights, participants completed each day the TloadDBack cognitive fatigue (CF) induction protocol under either high cognitive load (HCL) or low cognitive load (LCL) conditions, in a within-subject, counterbalanced design. Prior to, and immediately following the TloadDBack practice (TTask), participants’ EEG was recorded during a quiet resting period. Those pre-task (resting state 1 [RS1]), and post-task (resting state 2 [RS2]) periods spent under EEG had a duration of five minutes, in which participants were seated with the eyes open, and instructed to fixate a white cross on a black background. The 16-minutes TloadDback task (TTask) practice was also performed under EEG. Cognitive fatigue was subjectively evaluated using visual analog scales for fatigue severity (VASf; Lee et al.,1991) and sleepiness (VASs) at each experimental phase (RS1, TTask, RS2), alongside visual analog scales for stress (VASst) and motivation (VASm) to account for potential confounds. To mitigate interindividual circadian variability, each participant was assessed at the same time of day across all UdS and SF, LCL and HCL conditions.

**Figure 1.**
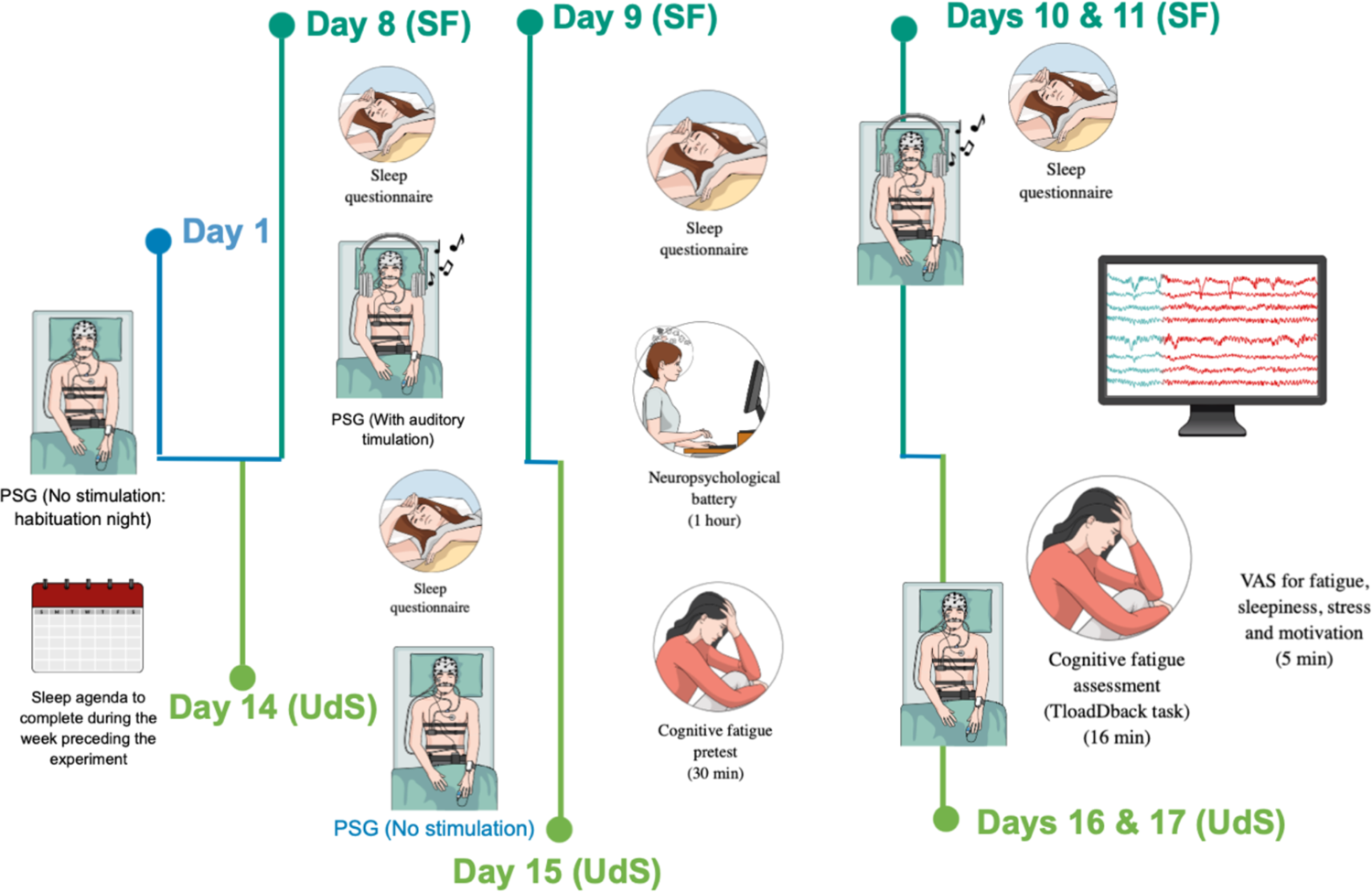
Experimental protocol (see general experimental procedure). Note: PSG = Polysomnography. VAS = Visual Analogue Scales (fatigue, sleepiness, stress, and motivation). LCL = Low Cognitive Load Condition. HCL = High Cognitive Load Condition. Participants spent three consecutive nights both in a sleep fragmentation (SF) and in a sleep undisturbed (UdS) condition at one-week interval, counterbalanced. Cognitive fatigue (CF) calibration (pretest) was performed after the first SF or UdS night, and CF assessment was conducted in HCL and LCL conditions after the second and third nights, counterbalanced.

**Figure 2.**
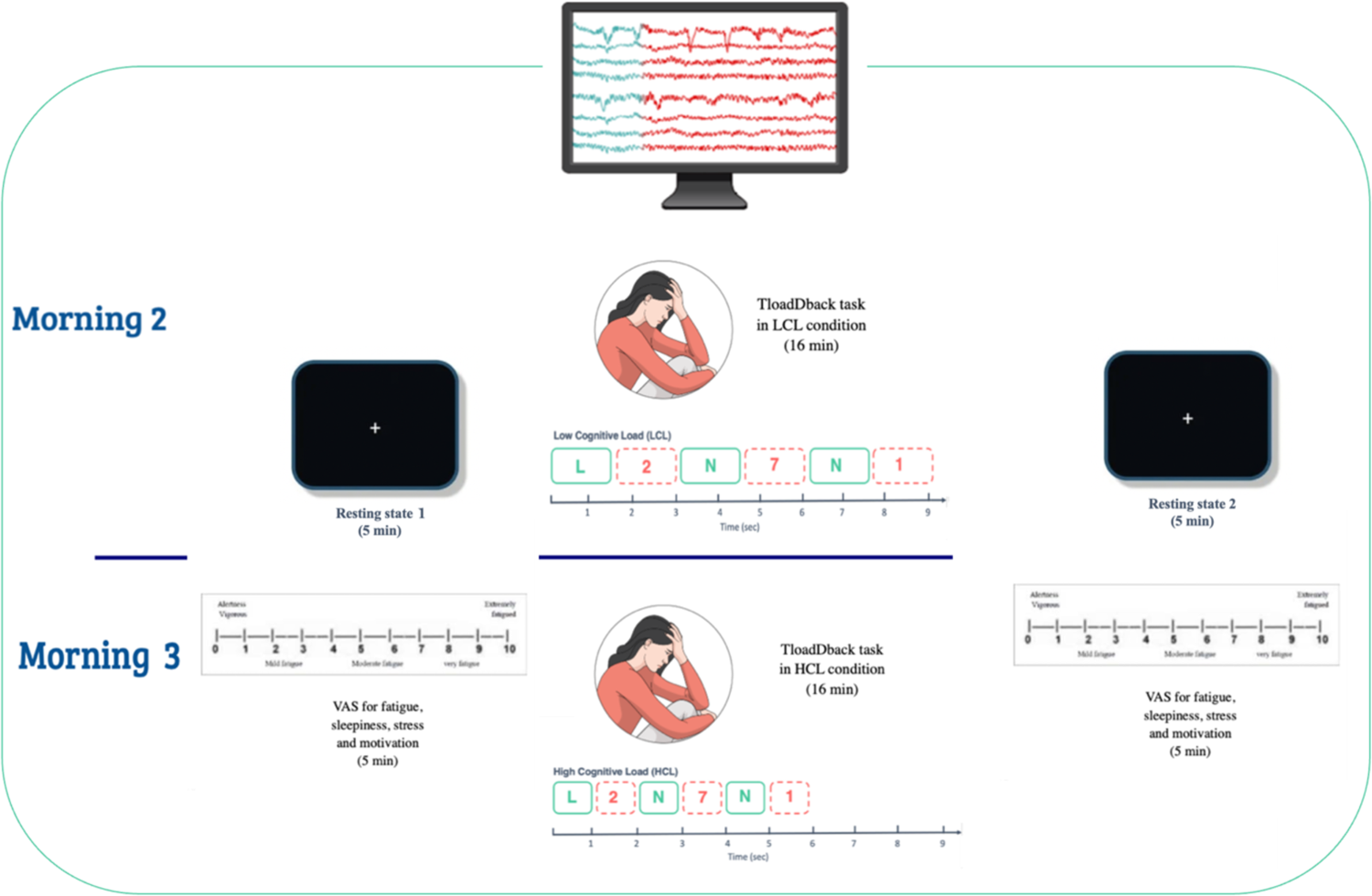
Timeline of resting states before and after the TloadDback task, performed under EEG. The TloadDback task was administered in both cognitive load conditions, counterbalanced, after SF/UdS nights 2 and 3.

### 2.3 Sleep Fragmentation procedure

Sleep fragmentation was induced using auditory stimulations at a frequency intended to simulate the sleep fragmentation experienced by patients with obstructive sleep apnea (for details, see Benkirane et al. 2022). The timing of the stimulations was personalized for each participant, considering their typical sleep duration. During each SF night, the initial sleep cycle remained undisturbed, after which arousing auditory tones were introduced through loudspeakers at random intervals ranging 60 to 120 seconds to induce SF. The auditory tones randomly alternated between beep tones and firecracker sounds to prevent habituation to the sounds or their repetition. The auditory stimulation started at a low intensity level and was gradually increased until microarousals were observed. If any signs of awakening were observed, as defined by the American Academy of Sleep Medicine (AASM) criteria, the generation of auditory tones was manually paused until the participant resumed a deeper stage of sleep (N2, N3, or REM) for at least 2 minutes, after which the SF protocol was resumed. In both SF and UdS nights, participants were informed that auditory tones might be presented without the intention of waking them up, but were not informed on which nights, or how many nights, it would occur. No information regarding the sleep condition was provided the following morning.

### 2.4 Cognitive Fatigue-Inducing TloadDback Task

The TloadDback task is a dual task that requires participants to perform an N-back working memory updating task and an odd/even number decision task simultaneously (Borragán et al., 2017). A detailed description of the task can be found elsewhere (Benkirane et al., 2022). In a nutshell, 30 digits and 30 letters are displayed on a screen in alternating order (e.g., N–2–X-7– X–1–L…), and participants must press the space key with their left hand every time the displayed letter is the same as the previous letter (1-back task; e.g., …X-7–X…), and indicate whether the displayed digit is odd or even by pressing the “2” or “3” keys with their right hand. The cognitive resources required for the task were individually adjusted during separate calibration sessions that took place after the first SF or UdS night (Days 9 and 15). Participants completed a maximum of 20 blocks, with an initially comfortable interstimulus interval (ISI) of 1500 ms for the first block. If the participant achieved an accuracy of at least 85% at a block, then the ISI for the next block was decreased by 100 ms, making the task more challenging by allowing less processing time. This staircase procedure was repeated until participant’s performance dropped below 85% over three consecutive blocks, indicating that they had reached their cognitive load limit. During the experimental conditions of SF or UdS (Days 10-11 and 16-17), participants completed the TloadDback task at a constant ISI for 16 minutes either in HCL or a LCL condition, counterbalanced. In the HCL condition, the ISI was set to the last successful ISI at the calibration session plus 100 ms. In the LCL condition, the ISI was one-third longer than in the HCL condition (ISI (LCL) = ISI (HCL) + 1/2 ISI (HCL)).

### 2.5 Electroencephalography

#### EEG recordings

EEG signals on the second and third day of each sleep condition (SF and RS) were recorded, amplified, and digitized at 256 Hz sampling rate with 24-bit resolution with the 32 channel SD LTM Express Morpheus system (Micromed, Mogliano Veneto, Italy) digital recorder operated with BrainRT software (OSG, Rumst, Belgium). PSG recordings included nine channels at scalp locations F4, Fz, F3, C4, Cz, C3, P3, Pz, and P4, referenced to the mastoids and placed according to the international 10-20 system (Keil et al., 2014), and bipolar EOG and EMG channels. The impedance of electrodes was kept below 5 kΩ. Participants’ skin was prepared according to standard procedures. All EEG recordings were exported as EDF+ files and preprocessed using the Fieldtrip toolbox (Version 20221223; Oostenveld et al, 2011) on MATLAB (Version R2021b; The MathWorks, Inc., Natick, MA). EEG data collection spanned 26 minutes, encompassing the 5 minutes prior to (RS1), the 16 minutes during (TTask), and the 5 minutes following (RS2) the completion of the TloadDback task.

#### EEG processing

Raw EEG data were bandpass filtered between 1 Hz and 30 Hz (with Butterworth, zero phase forward and reverse digital filter) to exclude low-frequency drifts and high-frequency noise. Independent component analysis (ICA) of the 5-minute long recordings of pre-and post-task resting states, as well as of data collected during the 16-minute long TloadDback task was performed to identify eye-movement and muscular artefacts using Fieldtrip routines (Oostenveld et al., 2011). ICA was performed on the concatenated pre-and post-task resting states, and on the recordings during the TloadDback task, separately. Independent components (mostly one, maximum two) representing components linked to eye movements and muscular artefacts were detected semi-automatically and were identified by inspecting the waveforms, as well as their topographical distribution (Campos Viola et al., 2009). The 16-minute TloadDback task was cut into 4 successive segments of 4 minutes each in order to investigate neurophysiological differences between the beginning and the end of the task, linked to the gradual development of CF during task practice. Recordings were further segmented into 4-sec long trials, visually inspected to exclude trials with remaining artifacts. Fast Fourier Transformation was performed on Hanning tapered, non-overlapping 4-sec long trials with 0.25 Hz resolution, to extract the averaged power spectral density of EEG activity in pre- and post-task resting states, as well as in the TloadDback task. Absolute power values were computed between 1.5 Hz and 25 Hz.

### 2.6 Statistics

Multiple comparisons were performed to contrast bin-wise absolute power between RS1 (pre-task) and RS2 (post-task) within the same sleep condition (SF and UdS) and within the same cognitive load condition (HCL and LCL) to test the fatigue effect linked to cognitive load and sleep fragmentation. The first and last 4-minute segments of each TloadDback condition were also compared across sleep (SF and UdS) and cognitive load (HCL and LCL) conditions. We expected that the HCL and SF conditions would induce changes in power spectral density (PSD). Differences between conditions were evaluated by non-parametric bootstrapping (Monte Carlo simulation with 1000 shuffled samples) (Maris & Oostenveld, 2007) as implemented in Fieldtrip (Oostenveld et al, 2011). Such approach is suitable for the analyses of EEG signals as it does not require assumptions of data distribution (Maris & Oostenveld, 2007). Two-tailed, paired t-tests were performed on all pairs of bin-wise absolute spectra (between 1.5-25 Hz) across two conditions. Spectral power values were averaged over the nine channels. To address the issue of multiple comparisons, spectral power differences along frequency ranges were considered as significant if they spanned along minimum three consecutive bins (with a threshold p <0.05), since there is a low probability to obtain significant differences in adjacent bins by chance (Tarokh et al., 2010). Significant bins were visualized, but were not considered significant if they did not reach the threshold. To explore and highlight the topographical aspects of spectral power differences, we also visualized significant differences in the frequency x channel space.

In order to ensure the integrity of the data analysis, participants with excessive noisy data were excluded from the study. Therefore, when analyzing the TloadDback task, data from 15 participants were kept in the HCL and UdS conditions, and in the LCL and SF conditions. Data from 14 participants were kept in the HCL and SF conditions, and in the LCL and UdS conditions. When analyzing resting states data, the 16 participants were kept in the HCL and SF conditions, and in the LCL and SF conditions. Data from 15 participants were kept in the HCL and UdS, and LCL and UdS conditions.

## 3. Results

### 3.1 Resting states EEG

The 5-minutes EEG data before (first resting state [RS1]) and after (second resting state [RS2]) the TloadDback task were analyzed within each condition (high(HCL)/low(LCL) cognitive load, and fragmented(SF)/undisturbed sleep(UdS). In case of high cognitive load conditions, significant differences between pre- and post-task resting states emerged in the theta (∼4.75-6 Hz), alpha (∼9-12.25 Hz) and beta frequency ranges (∼18-19 Hz) when participants performed the task after SF. More specifically, post-task resting EEG power increased compared to pre-task activity in these frequency ranges. Topographical analyses including all channels indicated widespread changes extending throughout the scalp (see Figure 3). In the case of undisturbed sleep (UdS), an increase in posterior low beta power (∼13-14 Hz) was observed from pre-to-post task resting state (after the high cognitive load task). No significant differences emerged between pre-and post-task resting states in the low cognitive load conditions, after sleep fragmentation or undisturbed sleep conditions either (See Figure 3).

**Figure 3.**
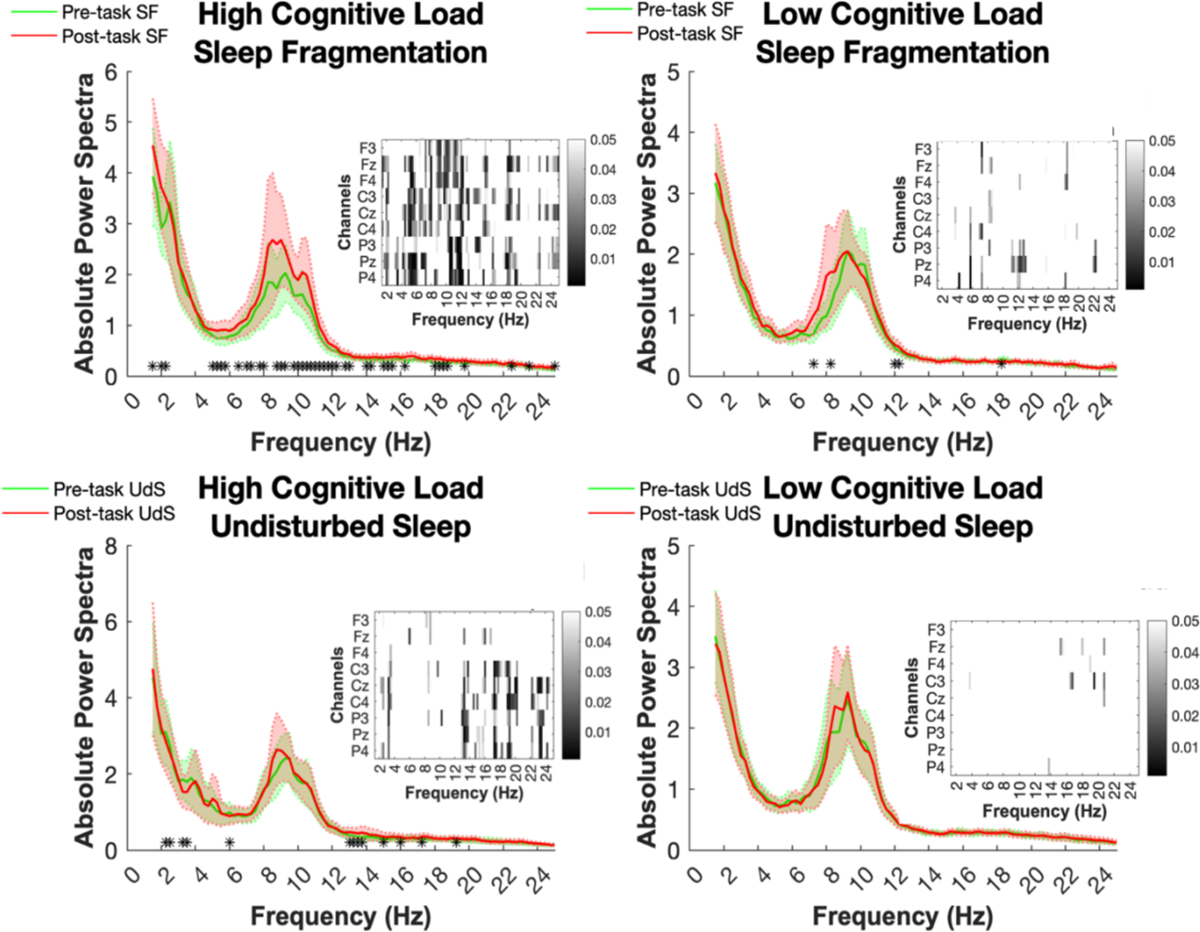
Visualization of Absolute Power Spectra during Resting States across contrasted conditions. Panels highlight the comparison between resting state EEG power before the TloadDback task (RS1) and after the TloadDback task (RS2) under high (HCL) and low (LCL) cognitive load conditions, under sleep fragmentation (SF) and undisturbed sleep (UdS) conditions. Inset heatmaps show channel-wise changes in the power spectra across frequencies for the specified conditions. Each heatmap displays the EEG channels (F3, F4, C3, C4, P3, P4, Cz) on the y-axis and frequency on the x-axis. Greyscale intensity indicates statistical significance of power change, with scale on the right-side axis (0 to 0.05). Significant differences are marked with * (p < 0.05). To address for multiple comparisons, spectral power differences along frequency ranges were considered significant only if they spanned along at least three adjacent bins.

A comparison between the first resting state (RS1) following SF vs. UdS conditions, with RS1 averaged across high and low cognitive load conditions, did not disclose any baseline difference, suggesting that identified differences at the second resting state (RS2) are linked to the developing cognitive fatigue (CF) during the intermediate TloadDback task practice.

### 3.2 TloadDback EEG

As a reminder, the 16-minutes EEG data collected in each condition (HCL/LCL, and SF/UdS) were segmented into four 4-minute sections. To assess the evolution of fatigue markers throughout task practice, the first and the last segments of the tasks were contrasted for each condition (HCL/LCL and SF/UdS). As shown Figure 4, we observed a significant difference in beta power (∼14-24 Hz) between the first and the last segment of the task in the high cognitive load and sleep fragmentation (HCL/SF) condition. More specifically, beta power exhibited a relative increase in the last segment of the task compared to the first segment. A similar increase in beta power, starting from the low beta range (∼12.5-24 Hz) was found in the last compared to the first segment of the task in the low cognitive load and undisturbed sleep (LCL/UdS) condition. No significant differences emerged in the other conditions (see Figure 4.)

**Figure 4.**
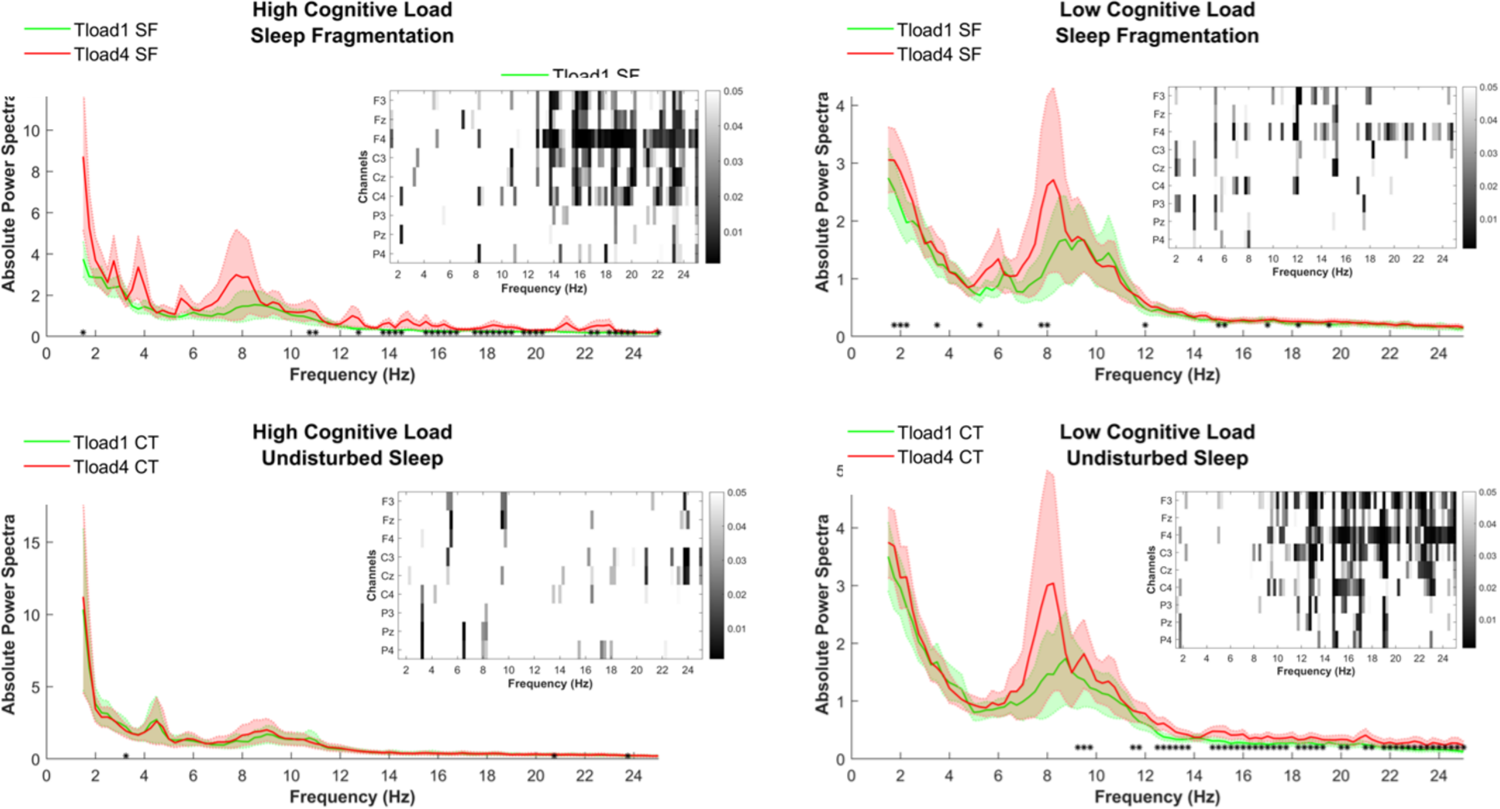
Visualization of Absolute Power Spectra during the first and last segment of the TloadDback task across conditions. Figure panels illustrate power spectra in the first 4-minutes segment (Tload1), and the last 4-minutes segment (Tload4) of the TloadDback task under different cognitive load (HCL and LCL) and sleep (SF and UdS) conditions. Inset heatmaps show channel-wise changes in the power spectra across frequencies for the specified conditions. Each heatmap displays the EEG channels (F3, F4, C3, C4, P3, P4, Cz) on the y-axis and frequency on the x-axis. Greyscale intensity indicates statistical significance of power change, with a legend on the right showing the scale (0 to 0.05). Significant differences are marked with * (p < 0.05); however, to address for multiple comparisons, spectral power differences along frequency ranges were considered as significant if they spanned along minimum three adjacent bins.

## 4. Discussion

In the present study, young healthy adults experienced three consecutive nights of auditory-induced sleep fragmentation (SF) and three nights of undisturbed sleep (UdS), in a counterbalanced design. We previously showed an effect of SF on sleep architecture, subjective appraisal of fatigue, and performance, even though total sleep time was equivalent between the two conditions. Noticeably, our young adult participants proved able to compensate for consequences of altered sleep continuity at the behavioral level in several cognitive domains (Benkirane et al., 2022). Here, we analyzed CF- and SF-related changes in EEG during the resting states before (RS1) and after (RS2), and during the fatigue inducing TloadDback task in high (HCL) and low (LCL) cognitive load conditions following either SF or UdS.

Beta activity in the resting state increased from before (RS1) to after (RS2) the TloadDback task in the high cognitive load (HCL) condition after sleep fragmentation (SF). Additionally, there was a notable rise in the high beta frequency band between the first and last segments of the TloadDback task in HCL/SF and the LCL/UdS conditions. These findings suggest that under conditions of high cognitive load combined with sleep fragmentation, the brain exhibits increased beta activity. Beta rhythms have been often associated with active concentration, alertness and top-down control mechanisms (Egner & Gruzelier, 2004; Stoll et al., 2016). They are also linked to attention to impending processes and their execution (Khanna & Carmena, 2015). Furthermore, beta rhythms have been found to encode information about task rules in the frontal cortex and are crucial for the activation of task sets in response to task demands (Spitzer & Haegens, 2017). This suggests that beta rhythm-related processes are essential to consider when examining the effects of fatigue during cognitive tasks, particularly those involving the initiation of motor processes following the presentation of target stimuli. Previous research has demonstrated a positive correlation between high-beta activity and the number of microsleep-associated errors, along with a decrease in hit ratio. This has been interpreted as an effort to stay awake rather than an indication of increased alertness, which interferes with performance and leads to an increase in task-related errors (Mairesse et al., 2009). Moreover, psychosocial stress induced by cognitive tasks has been shown to increase relative beta power during correct trials of an attentional task, correlating positively with anxiety and heart rate increase, and inversely with attentional accuracy (Palacios-García et al., 2021). In conditions of low cognitive load and undisturbed sleep conditions, the increase in both low and high beta activity over the task duration suggests that the brain still engages in heightened cognitive processing, possibly to maintain performance levels (Engel & Fries, 2010). These convergent results indicate a consistent pattern of beta band modulation in response to cognitive demands. Engagement in tasks of varying difficulty leads to an increase in beta oscillatory activity, highlighting the dynamic nature of beta rhythms in response to cognitive workload variations and sleep quality (Bonnet & Arand, 2003).

Additionally, we found a trend towards positive changes in power frequencies between the first and second 4-minute segments in the delta power band in the LCL/SF conditions. Delta oscillations, typically ranging from 0.5 to 4 Hz, are associated with deep sleep, and cognitive processing during sleep (Steriade et al., 1993). While the precise role of delta activity in cognitive fatigue is still under investigation, augmented delta power has been reported in conjunction with mental fatigue (Belyavin & Wright, 1987). The trend towards positive changes in delta power between the first and second 4-minute segments may indicate that the brain could allow some sleep-like activity in a less demanding task after SF. Durmer and Dinges (2005) showed that cognitive fatigue, often induced by sleep deprivation, can lead to microsleep episodes. These brief lapses in consciousness, lasting from a fraction of a second to several seconds, are characterized by increased delta activity and reduced responsiveness to external stimuli. This aligns with the subjective reports of higher baseline of subjective fatigue after SF but a larger increase following UdS (Benkirane et al., 2022), suggesting that the brain may be adopting a compensatory mechanism to cope with the accumulated fatigue, which needs to be confirmed in further studies.

It is only during the resting state following the more demanding cognitive task after sleep fragmentation (HCL/SF conditions) that alpha waves increased. This may indicate a rebound effect where the brain, after being taxed by high cognitive load and fragmented sleep, shifts toward a more relaxed state once the task is completed. This aligns with the theory that alpha oscillations are markers of relaxation and reduced sensory processing (Klimesch, 1999). Moreover, psychosocial stress linked to the demanding cognitive tasks has been shown to induce an increase alpha power synchronization (Palacios-García et al., 2021). Previous studies suggest that higher beta frequencies, along with alpha activity during eyes-open tasks, increase as a result of sleep resistance mechanisms (Mairesse et al., 2009). In the EEG signal, cognitive fatigue typically leads to an increase in parietal alpha power (Trejo et al., 2015), indicating a shift from active attention to the default mode, and a suppression of external stimulus processing (Klimesch, 1999; Hanslmayr et al., 2011). This rise in alpha activity is often paired with an elevation in frontal theta power, which indicates cognitive control, task execution, memory function, and error processing (Sauseng et al., 2010). Increasing theta activity is believed to indicate greater exertion to manage heightened cognitive demands during extended tasks. Furthermore, heightened power in slow frequency bands is linked to diminished performance, underscoring that prolonged mental fatigue can degrade task performance over time (Hamann & Carstengerdes, 2023). The trend observed in our study toward increased theta activity between the first and last segments of the TloadDback task in the HCL/SF condition may reflect a compensatory mechanism, as theta waves are not only indicative of working memory and cognitive load processing (Klimesch, 1999), but also suggest adaptive brain functioning. This increase in theta oscillations could signify an effort to maintain cognitive performance despite the challenges posed by sleep fragmentation and high cognitive load, potentially highlighting the brain’s adaptive response to mitigate the effects of cognitive fatigue.

Previous publications reported augmented delta and theta activity accompanying fatigue, alongside diminished beta activity concurrent with declining performance, vigilance and sustained attention (Belyavin & Wright, 1987; Klimesch, 1999; Makeig et al., 2000; Oken et al., 2006, Lal & Craig, 2002). It suggests that increased delta and theta oscillations may reflect the neurophysiological manifestations of cognitive fatigue. Changes in delta power also align with behavioral outcomes indicating a decline in performance over practice time, emphasizing the complex relationship between neural markers and cognitive fatigue manifestations (Gorgoni et al., 2020). In studies examining EEG patterns, consistent findings include increases in delta and theta rhythms as drowsiness and fatigue build up, especially in frontal and central areas. Increased theta activity, in particular, has been correlated with decrements in cognitive task performance (Hamann & Carstengerdes, 2023). EEG activity has been shown to be highly sensitive to cognitive fatigue over extended periods, characterized by substantial increases in frontal theta and parietal alpha power, even preceding noticeable performance decline. Our CF-inducing task, although relatively short (16 minutes), has been carefully designed to induce cognitive fatigue effectively. This design was informed by pretest calibrations tailored to each participant’s specific cognitive abilities, ensuring that the task reliably induces fatigue within the given timeframe.

In line with other studies (Cote et al., 2003; Kingshott et al., 2000; Ko et al., 2015), our previous research demonstrated that sleep fragmentation, while maintaining total sleep duration, minimally impacts behavioral performance across various cognitive domains (Benkirane et al., 2022). However, neurophysiological measures reveal heightened sensitivity to even moderate sleep disruptions (Ko et al., 2015). The current findings underscore the significant impact of cognitive load and sleep quality on neural activity patterns, highlighting the intricate interplay between cognitive demands, brain oscillations, and cognitive fatigue (Cano et al., 2008). Specifically, alpha and theta rhythms appear pivotal in the context of cognitive fatigue and task performance, suggesting their potential as neural markers for monitoring mental workload and fatigue levels (Yu et al., 2021). Moreover, increases in parietal beta power have been associated with emerging mental fatigue (Dasari et al., 2010; Nguyen et al., 2017). Tanaka et al. (2012) propose that the observed enhancements in beta and alpha power densities during mental fatigue may signify a decline in multi-modal, high-level information processing within the central nervous system.

In contrast to the significant changes observed under the high cognitive load (HCL) and sleep fragmentation (SF) conditions, we did not observe significant changes in alpha, beta, or theta power under the low cognitive load (LCL) conditions following sleep fragmentation (SF), and high cognitive load (HCL) conditions following undisturbed sleep (UdS), suggesting that the interplay between cognitive load and sleep quality may differentially impact neural activity patterns. The lack of observable changes in these conditions may indicate that compensatory brain mechanisms are more effective when either cognitive load is lower or sleep quality is higher, allowing to maintain a more stable EEG activity. This stability may reflect an adaptive response, where the brain is better able to cope with cognitive demands under less stressful conditions, or when undisturbed sleep has mitigated the adverse effects of sleep fragmentation. It highlights the importance of considering both cognitive load and sleep quality together, as their interaction appears to play a crucial role in influencing neural activity and cognitive performance.

## 5. Limitations

While our study can offer valuable insights into the impact of sleep fragmentation on cognitive fatigue and its neurophysiological correlates, several limitations warrant consideration. Our population sample composed of young and healthy adult may restrict the generalizability of our findings to broader populations, especially in pathological contexts. Additionally, the experimental induction of sleep fragmentation may not fully replicate real-world sleep disturbances that usually span over much longer periods of time, potentially limiting the ecological validity of our results. Finally, the cognitive load task design and EEG measurements, while carefully constructed, may not fully capture the complexity of everyday cognitive demands and neural responses. Despite these limitations, our study provides a foundation for future research to explore these phenomena in more diverse and ecologically valid contexts.

## 6. Conclusions

Our EEG study examined the effects of sleep quality and cognitive load on neurophysiological markers of cognitive fatigue in young adults. Significant alterations in EEG activity were observed under high cognitive load (HCL) and sleep fragmentation (SF) conditions, with increased beta and alpha activities indicating heightened cognitive effort and relaxation rebound, respectively. A trend for increased theta activity suggested compensatory mechanisms to maintain performance. In the low cognitive load (LCL) and undisturbed sleep (UdS) conditions, increases in low beta power density suggest that the brain engages in heightened cognitive processing even under less demanding circumstances to maintain performance levels. These findings underscore the roles of alpha and theta rhythms in cognitive fatigue and task performance, suggesting their potential as neural markers for monitoring mental workload. Overall, the study highlights the interplay between sleep quality and cognitive load, emphasizing their combined impact on cognitive performance and neural activity. These insights may contribute to understanding cognitive fatigue mechanisms and offer implications for managing fatigue in conditions like sleep apnea.

